# Fishing for contact: Modeling perivascular glioma invasion in the zebrafish brain

**DOI:** 10.1101/2020.08.15.252544

**Authors:** Robyn A. Umans, Mattie ten Kate, Carolyn Pollock, Harald Sontheimer

**Affiliations:** Center for Glial Biology in Health, Disease, and Cancer, The Fralin Biomedical Research Institute at VTC, Roanoke, VA, 24016, USA; School of Neuroscience, Sandy Hall, 210 Drillfield Dr.,Virginia Tech, Blacksburg, VA, 24061, USA

**Keywords:** Glioblastoma multiforme, perivascular glioma invasion, zebrafish, tumor model, Wnt signaling

## Abstract

Glioblastoma multiforme (GBM) is a highly invasive, central nervous system (CNS) cancer for which there is no a cure. Invading tumor cells evade treatment, limiting the efficacy of the current standard of care regimen. Understanding the underlying invasive behaviors that support tumor growth may allow for generation of novel GBM therapies. Zebrafish (*Danio rerio*) are attractive for genetics and live imaging, and have in recent years, emerged as a model system suitable for cancer biology research. While other groups have studied CNS tumors using zebrafish, few have concentrated on the invasive behaviors supporting the development of these diseases. Previous studies demonstrated that one of the main mechanisms of GBM invasion is perivascular invasion, i.e. single tumor cell migration along blood vessels. Here, we characterize phenotypes, methodology, and potential therapeutic avenues for utilizing zebrafish to model perivascular GBM invasion. Using patient derived xenolines or an adherent cell line, we demonstrate tumor expansion within the zebrafish brain. Within 24 hours post-intracranial injection, D54-MG-tdTomato glioma cells produce finger-like projections along the zebrafish brain vasculature. As few as 25 GBM cells were sufficient to promote single cell vessel co-option. Of note, these tumor-vessel interactions are CNS specific, and do not occur on pre-existing blood vessels when injected into the animal’s peripheral tissue. Tumor-vessel interactions increase over time and can be pharmacologically disrupted through inhibition of Wnt signaling. Therefore, zebrafish serve as a favorable model system to study perivascular glioma invasion, one of the deadly characteristics that make GBM so difficult to treat.

---

Brain tumors are one of the most devastating forms of cancer, as they grow within an organ of limited regenerative capacity and can be highly invasive. Additionally, drug delivery to brain tumors is challenging due to the blood-brain barrier (BBB), a physiological structure that limits the open flow of foreign compounds into the brain tissue. Glioblastoma multiforme (GBM) is the most common, primary malignant brain tumor and is newly diagnosed in around 13,000 patients every year in the United States ^1^. The standard of care is a three-pronged approach of surgical resection, radiation, and chemotherapy ^2^. Even after over a decade of implementation, this multifaceted treatment regimen extends survival to only 14-19 months ^3^. These aggressive therapies mainly target the main tumor mass that is most easily identified by MRI for surgical removal or therapeutics that target actively dividing cells. Even in the era of personalized medicine, every GBM case presents with a different set of mutations, making it difficult to standardize a therapeutic regimen that effectively treats all patients. Research efforts have shown that the genetic heterogeneity of GBM tumors results in varied patient response to specific therapies, and individual mutations may not be present in the majority of primary GBM patients ^4–7^. Therefore, targeting how these brain tumor cells behave in the context of their mutations or environment may elucidate avenues for more promising therapies that have a wider reach within the GBM patient population ^8^.

One of the main routes of GBM infiltration is cellular invasion within the tight spaces of the brain parenchyma. While the main neoplasm is targeted for surgical resection, migratory cells can remain undetected by MRI and form satellite tumors away from the original mass. The avenues by which these motile cells travel are white matter tracts and blood vessels, with the perivascular space presenting little physical resistance ^9, 10^. Previous work using a mouse model intracranially injected with human GBM cells demonstrated human tumor cell tropism along the mouse brain vasculature, and after 2-4 weeks of implantation, GBM cells caused micro-breaches at the BBB ^11^. Furthermore, another GBM rodent model demonstrated the Wnt signaling cascade is necessary for glioma-vessel interactions ^12^. However, *in vivo* mouse models require the injection of thousands of cells, use of immune-compromised animals, and performing advanced surgical and imaging techniques. The emergence of an *in vivo* model that could more easily visualize and manipulate perivascular interactions important for tumor invasion would greatly benefit the field of GBM biology. In recent years, groups have established zebrafish (*Danio rerio*) specifically modeling the growth and invasion of brain tumors ^13–15^. These studies take different approaches, such as injecting brain tumor cells into the yolk of the animal or a large bolus into the brain to study invasion and tumor angiogenesis ^14, 16, 17^. While these zebrafish models provide exciting avenues to study human disease in a simpler vertebrate system, most have looked at the molecular features of tumors and not the mechanism of perivascular glioma invasion. Therefore, a more thorough characterization of the relationship between glioma cells with the zebrafish brain vasculature remains.

In the present study, we hypothesize that zebrafish could be utilized to specifically model perivascular glioma invasion. To test and visualize this hypothesis, we intracranially injected red fluorescently labeled human glioma cells into the translucent zebrafish vascular reporter line *Tg(fli1a:eGFP)^y1^;casper*, whereby all blood vessels are labeled with enhanced green fluorescent protein (eGFP). We injected both an adult glioma cell line (D54-MG) and a pediatric patient derived xenoline (D2159MG) into *Tg(fli1a:eGFP)^y1^;casper* zebrafish and witnessed both of these glioma lines attach to the zebrafish vasculature. Gliomas retain salient disease features in the zebrafish brain such as attachment to secondary structures (blood vessels). Injected animals were imaged daily and confocal microscopy revealed glioma cells survive and expand within the zebrafish brain. Perivascular glioma invasion was dependent on the number of tumor cells injected. When ≤25 cells were initially injected, tumors survived, grew, and had more contacts with the vasculature than if an animal was implanted with >25 cells. Tumor cell vessel co-option increased over time along the vessel and at vessel branch points. Glioma-vessel interactions were also dependent on the CNS microenvironment, as glioma cells injected into the animal’s peripheral tissue failed to interact with pre-existing vasculature. Furthermore, since prior studies have alluded to Wnt signaling being important in vessel co-option in a rodent glioma model, we examined whether requirement for Wnt signaling is conserved in zebrafish^12^. We utilize the small molecule XAV939, an inhibitor of the Wnt signaling pathway, to pharmacologically disrupt glioma cell-vascular interactions in zebrafish brain as it has been demonstrated in the mammalian brain. These proof-of-principle studies demonstrate valuable features of this established zebrafish model of perivascular glioma invasion, which could lead to the discovery of newly needed GBM therapeutics.

## Methods

### Cell Culture

D54-MG human glioma adherent cell lines (WHO IV, glioblastoma multiforme; gifted by Dr. D. Bigner, Duke University, Durham, NC) were genetically modified to express tdTomato or eGFP in a previous study ^11^. D54-MG cells were maintained in Dulbecco’s modified Eagle’s Media (DMEM) (Fisher #11320082) supplemented with 7% fetal bovine serum (Lot 105313, Aleken), and kept in a cell culture incubator with 10% CO_2_ at 37°C

D2159MG and GBM22 patient-derived xenograft (PDX) lines (gifted by Darell D. Bigner, MD, PhD and Stephen T. Keir, DrPH, MPH, Duke University, Tisch Brain Tumor Center and Dr. Yancey Gillespie, University of Alabama at Birmingham, Birmingham, AL, USA, respectively) were from primary brain tissue and maintained by serial passage in the flank of athymic nude mice, as previously described ^11^. PDX cells were grown in a DMEM complete media supplemented with B-27 (Fisher #12587010), Amphotericin (1:100; Fisher #BP264550), Sodium pyruvate (1:100;ThermoFisher #11360-070), EGF and FGF growth factors (10ng/ml each, final;ThermoFisher #PHG0266 and #PHG0315), and Gentamicin (1:1,000;Fisher #BW17 518Z). PDX cells were maintained in a cell culture incubator with 10% CO_2_ at 37°C. The D2159MG pediatric glioma cells were genetically modified with a lentivirus (MGH Vector Core) to constitutively express mCherry. To visualize unlabeled GBM22 cells, cells were labeled overnight with 5µM of the lipophilic DiL dye (Invitrogen #V22885).

For cell counting, D54-MG cells were dissociated using Trypsin-EDTA (ThermoFisher/LifeTech #25300054) and PDX cells were dissociated using Accutase (Sigma #A6964). Cells were counted on a hemocytometer and re-suspended at 25,000cells per µL in sterile PBS prior to microinjection.

### Acute Mouse Brain Slice Invasion and Immunohistochemistry (IHC)

Experiments were performed in accordance with the University of Alabama, Birmingham Institutional Animal Care and Use Committee (IACUC). *In situ* tumor invasion was performed as previously described ^18^. For IHC, 300µM brain sections were acutely sliced from 10 week old *NG2:dsRed* mice and seeded with 80,000 D54-MG-eGFP tumor cells on multi-well filter chambers (Fisher #08-771). Tumor cells invaded brain slices for 4 hours on filter chambers in a 10% CO_2_ cell culture incubator. Brain slices were fixed in 4% PFA and used for subsequent IHC. Slices were rinsed in PBST for 15 minutes, permeated with proteinase K (20mg/ml stock, 1:800) for 15 minutes at room temperature, and then blocked in 10% goat serum for 1 hour at room temperature. Sections were stained with a mouse-anti-Laminin primary antibody (Sigma L8271,1:500) overnight at 4ºC. The next morning, slices were washed in PBST for 1.5 hours, stained with donkey-anti-mouse-Alexa 555 secondary antibody (Invitrogen A31570,1:500) for 3 hours at room temperature, washed in PBST for 1.5 hours, and mounted with Fluoromount (VWR 100502-406) on glass slides. Z-stacks were acquired with an Olympus Fluoview FV1000 confocal microscope and a 60X(NA1.42) oil objective.

### Zebrafish

Adult zebrafish were maintained according to Virginia Tech IACUC guidelines on a 14/10 hour light/dark cycle in a Tecniplast system at 28.5°C, pH 7.0, and 1000 conductivity. The *Tg(fli1a:eGFP)^y1^* line was crossed onto the *casper* background ^19^ to generate genetically translucent vascular reporter lines. These animals were specific-pathogen free (SPF) and came from the Sinnhuber Aquatic Research Laboratory at Oregon State University. SPF animals were used for these brain tumor studies to avoid confounding factors such as *Pseudoloma neuropihila*, a pathogen that resides in the zebrafish CNS and is common in zebrafish colonies. The *Tg(glut1b:mCherry)* line was obtained from Dr. Michael Taylor’s laboratory at the University of Wisconsin-Madison. For all experiments, embryos were collected from multiple pair-wise crosses. Embryos were maintained in embryo water (1.5g Instant ocean salt per 5L RO water) at 28.5°C. For studies with the *Tg(glut1b:mCherry)* line, 24 hour post-fertilization embryos were treated with 200µM N-Phenylthiourea (Sigma # P7629) to prevent melanocyte formation. Embryos for tumor cell injections were gradually acclimated to develop at 32°C, so as to accommodate a more desirable temperature for glioma cell growth.

### Tumor Microinjections

Healthy animals (i.e. no developmental abnormalities) were screened before tumor implantation. 3 days post fertilization (dpf) *Tg(fli1a:eGFP)^y1^;casper* or *Tg(glut1b:mCherry)* larvae were anesthetized with 0.04% MS-222 (Sigma #A5040) prior to microinjection. During anesthesia, thin wall glass capillary needles with filament (World Precision Instruments #TW150F-4) were pulled with program 4 “Pro-Nuclear Injection” (Heat: 460, Pull:90, Vel: 70, Delay: 70, Pressure: 200, Ramp: 485) on a horizontal pipette puller (Sutter, Model P-1000). This program creates needles with a 0.7µM tip and a taper of 6-7mm. 1µL of Phenol Red (Sigma #P0290) was added to 9µL of the 25,000 glioma cell/µL solution mixture. Capillaries were positioned in a round glass pipette holder and back loaded with 1uL of human glioma cell mixture. Each loaded needle was calibrated using a pneumatic pump and micrometer (Carolina Biological Supply #591430) to calculate the bolus and corresponding cell number per injection. Animals were positioned dorsal side up for intracranial injection or on their sides for trunk injection in a homemade agarose injection mold, and then microinjected with the desired amount of 25 cells. While we attempted to only implant 25-50 cells (1nL) per animal, microinjections are not trivial and slight pressure differences between animals could lead to more or less cells implanted. After injections, were tracked in individual wells of a 24-well plate. After tumor microinjection, animals recovered for at least one hour, were anesthetized, and then embedded for live confocal microscopy to assess initial tumor volume.

### Live Confocal Microscopy

Animals were removed from multi-well plates with a glass Pasteur pipette and anesthetized with 0.04% MS-222 in individual 35mm petri dishes containing a 14mm glass coverslip (MatTek Corporation #P35G-1.5-14-C). A 1.2% low melting point agarose (ThermoFisher #16520050) solution was made in embryo water for embedding. MS-222 was removed from the animal and a bolus of warm agarose the size of the coverslip was added to the animal/cover glass. A dissection probe was used to align animals either dorsal side down for brain imaging or sagittally for trunk imaging. After the agarose solidified, the petri dish was filled with 0.04% MS-222 to keep the animal immobilized and the dish was wrapped with parafilm along the edges. Live confocal microscopy was performed to take z-stacks at optimal section thickness of animals with D54-MG and D2159MG cells with an Olympus Fluoview FV1000 confocal microscope and a 40X(NA0.75) objective. The Nyquist optimal section thickness value for z-stacks was calculated through an algorithm in the Olympus Fluoview software for all images. For whole zebrafish brain imaging after GBM22 injections, a Nikon A1R confocal microscope with a 10X(NA 0.45) objective and 1.5 optical zoom was utilized. Because these microscopes are upright, the glass coverslip bottom dishes containing the animals were flipped upside down so that the glass side made contact with the objective. Supplemental Movie 1 was generated from a Maximum Projection Image in NIS-Elements software.

### Small Molecule Treatment

After initial tumor implantation at 3dpf and subsequent imaging, tumors grew overnight before animals were unbiasedly sorted into treatment groups at 4dpf. The Wnt signaling inhibitor, XAV939, (Cayman Chemical #13596) was solubilized in dimethyl sulfoxide (DMSO) to generate 10mM stock solutions and used at a final concentration of 30µM in zebrafish egg water.

### Glioma-Vessel Interaction Analysis

To assess the percentage of human glioma cells interacting with the vasculature, OIB z-stack files were analyzed in the Fluoview software. In brief, the total number of visible human glioma cells were initially counted. The orthogonal view option was used to detect whether the signal from a cell was associated with the signal from a blood vessel or blood vessel branch point as seen in both XZ and YZ planes. The total number of interacting cells on a blood vessel or cells specifically at a vessel branch point was counted and then this value was divided by the total number of cells present at that point in time.

### *Tg(glut1b:mCherry)* Volumetric Analysis

Fiji-ImageJ and NIS-Elements software were utilized to measure the volume and signal intensity of *glut1* in tumor-associated and non-tumor associated vessels. In brief, confocal z-stacks were opened as split channels in FIJI and tiff files were saved to import into NIS-Elements. After importing, channels were merged together, the document was calibrated, and files were saved as a ND2 file. Sections around vessels of interest were cropped out of the whole file and a line was drawn along a vessel where it was ensheathed by a tumor cell body. The rotating rectangle feature was utilized and this selected area underwent a binary threshold and 1X clean and smooth options. The same procedure and size line was drawn along a similarly placed vessel without a tumor cell in the contralateral brain as a paired control vessel, to measure the volume and intensity in a similar manner. Measurements were generated in the NIS Elements “3D Object Measurements” table.

### Statistical Analysis

For analysis of perivascular glioma invasion at developmental time points 3-7dpf, a one-way ANOVA with multiple comparisons test was performed using GraphPad Prism software to determine *p*-values. For small molecule treatment experiments, a two-tailed, unpaired student’s *t*-test was performed using GraphPad Prism software to determine *p*-values. For volumetric vessel and *glut1* fluorescence analyses, a two-tailed, paired student’s *t*-test was performed using GraphPad Prism software to determine *p*-values.

## Results

### Human tumors retain salient features in a cell number-dependent manner in the zebrafish brain

While traditional glioma models utilize rodents studies have now demonstrated zebrafish are a suitable model to elucidate CNS cancer biology^16, 17, 20^. As a means to ask whether glioma-vessel interactions may occur in the zebrafish brain, we opted to inject fewer cells than previously published. Animals from the optically translucent *Tg(fli1a:eGFP)^y1^;casper* vascular reporter line were injected with 25-50 human D54-MG-tdTomato cells at 3 dpf, a time point when the BBB has started forming and sufficient CNS angiogenesis has occurred ^21^. Similar to the behavior of the D54-MG tumor line in acute mouse brain slices (**Figure 1A**), D54-MG glioma cells attached to the zebrafish brain vasculature within 24 hours post-intracranial injection (**Figure 1B,C**). We also performed microinjection into non-SPF animals and witnessed glioma-vascular interactions (**Supplemental Figure 1**), suggesting that while maintaining a SPF colony is the best practice for studying diseases of the CNS, it does not affect the growth of human brain cancer cells with regard to the zebrafish brain vasculature. This data suggests conserved signaling mechanisms exist to promote the interaction between glioma cells and the developing zebrafish vessel network.

**Figure 1.**
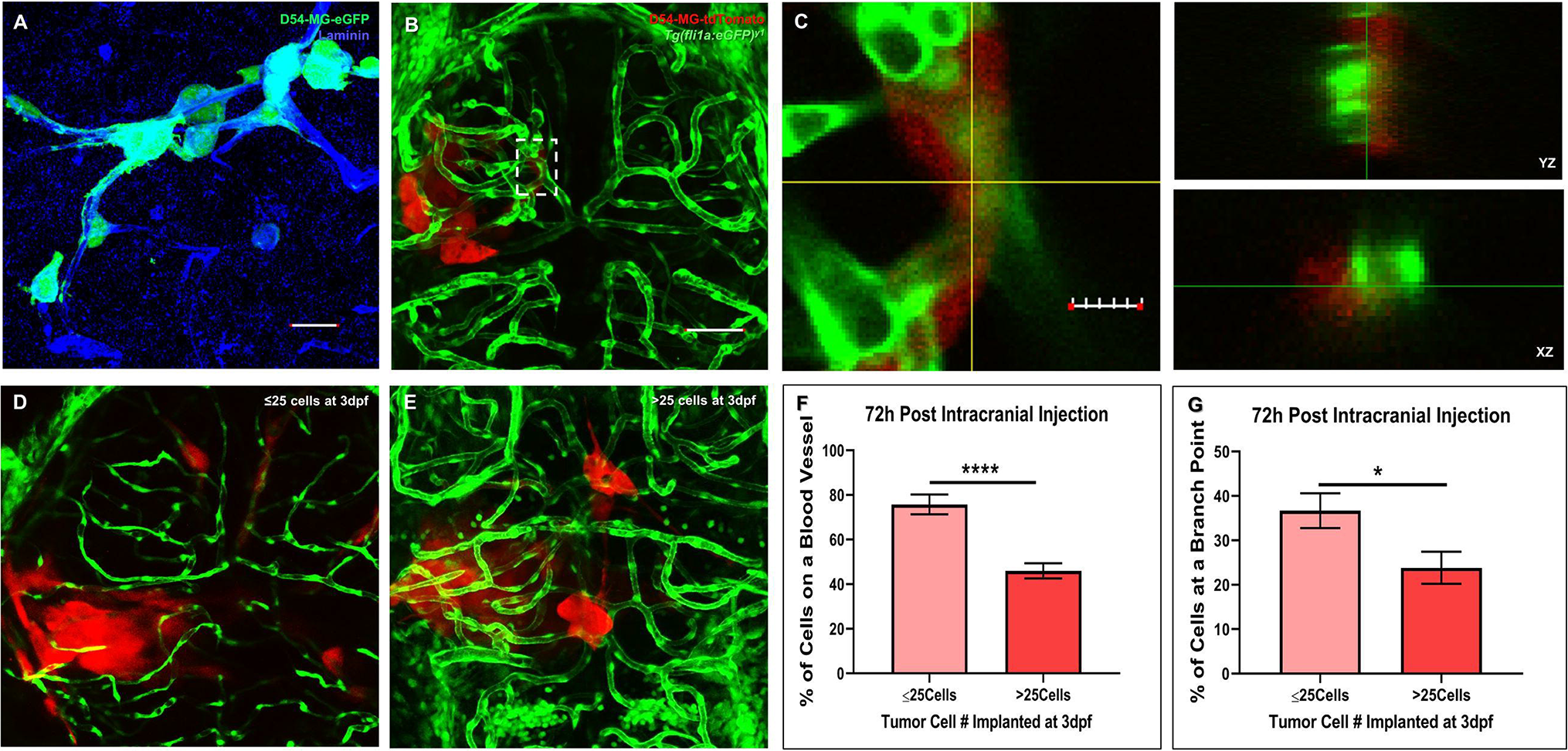
Glioma cells retain salient features in the zebrafish brain. (A) A maximum intensity projection of acute slice invasion with mouse brain seeded with 80,000 D54-MG-eGFP tumor cells. After 4 hours of invasion, slices were fixed and stained for the vessel protein Laminin (blue) to visualize the close association of glioma cells (green) during perivascular glioma invasion in this acute *ex vivo* model. Scale bar= 25µm. (B) The transgenic zebrafish vascular reporter line *Tg(fli1a:eGFP)^y1^;casper* (green) was intracranially injected with 25-50 D54-MG-tdTomato glioma cells (red) at the midbrain-hindbrain boundary at 3 days post-fertilization (dpf). 24h post-injection, a maximum intensity projection of an injected animal reveals tumor-vessel associations as seen in the acute slice model. Scale bar= 50µm. (C) An area (dotted white box in B) of a representative Z-plane from the confocal stack showing the corresponding orthogonal planes between the vessel (green) and glioma cell (red) signals. Scale bar 10µm. (D,E) Representative confocal images of 6dpf animals initially injected with ≤25 cells (D) or >25 cells. (F,G) Representative live confocal images of *Tg(fli1:eGFP)^y1^* larvae (red), 72h post-injection with ≤25 cells (D) or >25 cells (E). (F,G) Quantification of tumor-vessel interactions after 72h post-injection, comparing animals implanted on 3dpf with either ≤25 cells or >25 cells. Data shows tumor-vessel interactions along the vessel wall (F) or at a branch point (G) (n=. A two-tailed, unpaired student’s t-test was performed. Error bars represent mean with standard error of the mean(**p=<0.01), *p=<0.05). (n=16-21 animals).

To decipher whether glioma-vascular interactions depended on the volume of implanted cells, we intracranially injected a cell bolus of 25-50 and ≥50 glioma cells in 3dpf *Tg(fli1a:eGFP)^y1^;casper* animals. We observed that a larger bolus of ≥50 cells became more easily trapped within the ventricular space (data not shown), which is superficial to the developing zebrafish brain and therefore why we decided to perform the remaining experiments with lower injection volumes. These >50 cell compacted tumors were reminiscent of previously published zebrafish glioma studies that, due to cell number and size of the zebrafish brain versus human cells, could not examine the specific behaviors associated with perivascular glioma invasion ^16^. However, when 25-50 cells were injected into the midbrain-hindbrain boundary at 3dpf, we witnessed frequent extension of finger-like processes as previously observed in GBM mouse models and human brain ^11^. While present, >25 cell injected animals had fewer tumor-vessels interactions, along the vessel wall or at a branchpoint, within a 72 hour timespan compared to animals implanted with ≤25 cells at 3dpf (**Figure1D-G**). While survival studies were not in the scope of this study, some animals would succumb to the disease 4 days post-injection (data not shown). Previous studies report a median survival time of 10 days post injection in zebrafish with glioma xenografts of the preferred cell number (~25-50 cells) used in our study ^14^. Taken together, these experiments reveal that salient features of GBM and the process of perivascular invasion both occur in the developing brain of our zebrafish glioma model.

### Glioma cells actively invade the brain along the zebrafish vasculature

To analyze the dynamics of perivascular glioma invasion in the developing zebrafish brain, we microinjected D54-MG-tdTomato tumor cells into the midbrain-hindbrain boundary of 3dpf *Tg(fli1a:eGFP)^y1^;casper* larvae and monitored tumor behavior until 7dpf (**Figure 2A**). Because daily live imaging is easy to perform with the zebrafish model system, we wanted to assess tumor burden after the onset of injection and subsequent days. As shown by representative images in **Figure 2**, the few D54-MG-tdTomato glioma cells implanted at 3dpf (**Figure 2B**) not only survived in the zebrafish brain, but also expanded and migrated along the vascular network by 7dpf (**Figure 2C**). These glioma-vessel interactions increased over time along the main vessel wall (**Figure 2D**) and at branch points (**Figure 2E**) as previously described in a GBM mouse model ^11^. Furthermore, these interactions occurred with both pediatric and adult glioma cells, as well as in adherent and PDX cell lines (**Figure 3**). With a variety of PDX lines at our disposal, we labeled the GBM22 PDX line with a lipophilic dye to visualize tumor cells post-intracranial injection. Of note, lipophilic dye did not permeate the cell’s finer processes after injection and therefore does not retain the morphology that was visualized with D54-MG-tdTomato cells (**Supplemental Figure 2).** Therefore, it was determined that tumor cells that are transgenically labelled to express a fluorophore are more suitable for these imaging studies. These results demonstrate that even though glioma injections occurred in the developing brain, conserved signaling pathways support perivascular glioma invasion over time in our zebrafish model.

**Figure 2.**
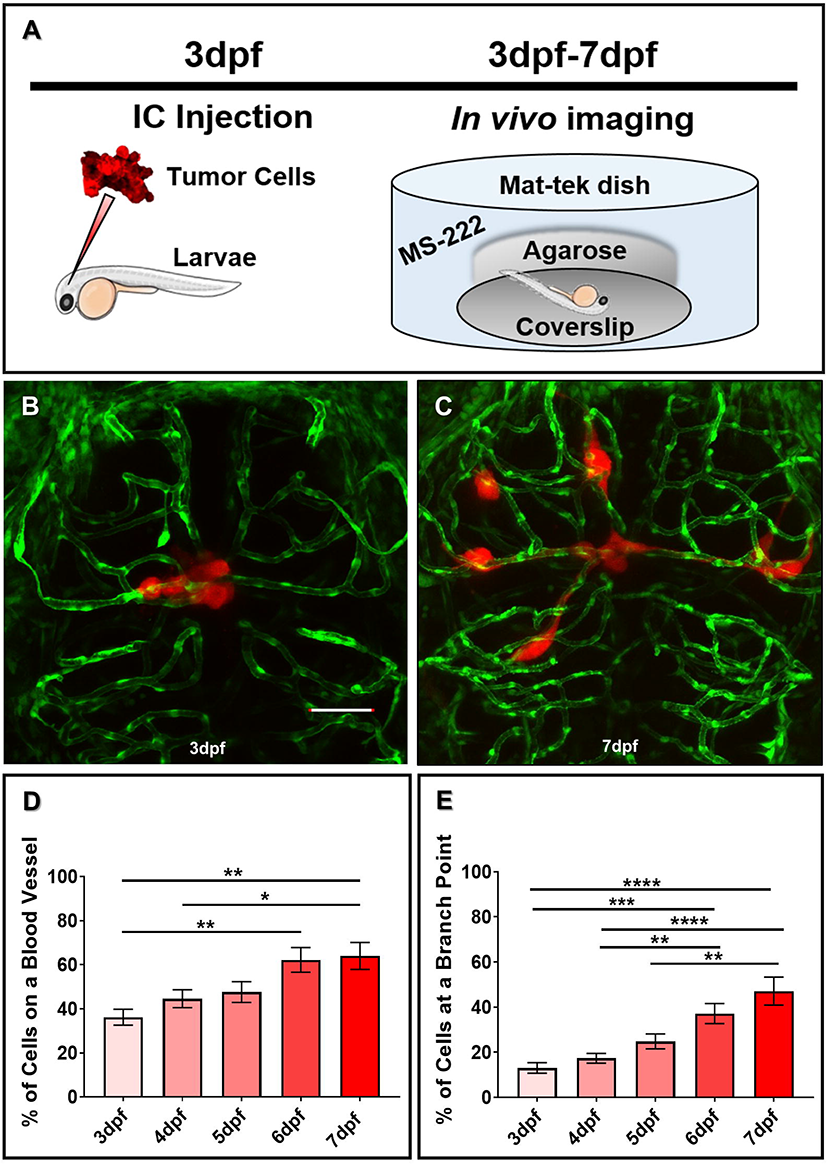
Glioma cells actively associate with the developing zebrafish brain over time. (A) A cartoon schematic of intracranial (IC) injection and subsequent *in vivo* imaging of our zebrafish perivascular glioma model. *Tg(fli1a:eGFP)^y1^;casper* animals were injected with 25-50 glioma cells in the midbrain-hindbrain boundary at 3dpf. After recovering, animals were imaged every day until 7dpf to monitor tumor burden over time. (B,C) Representative maximum intensity projection confocal images of tumor cells implanted at 3dpf (B) and subsequent migration by 7dpf (C). Scale bar= 50µm. (D,E) Quantification of tumor-vessel interactions in the developing brain. (D) Quantification of D54-MG-tdTomato cells attached to blood vessels over time (**p<0.001,*p=<0.05). (E) Quantification of D54-MG-tdTomato cells at vessel branch points over time (****p=<0.0001,***p=<0.001,**p=<0.01). *n*= 18-21 animals per time point. A one-way ANOVA with Tukey’s Multiple comparisons test was performed. Error bars represent mean with standard error of the mean.

**Figure 3.**
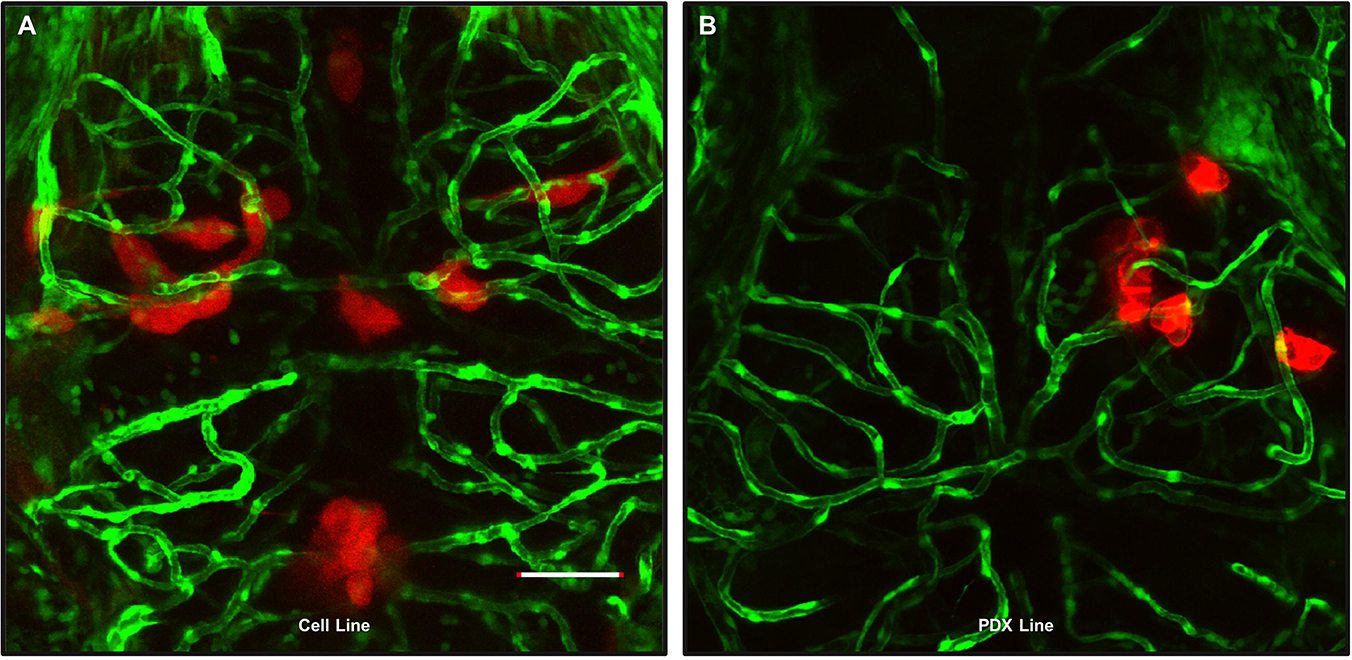
Glioma-vascular interactions occur in the zebrafish brain with various GBM lines. An adult D54-MG-tdTomato cell line (A, red) and D2159MG pediatric xenoline (B, red) were intracranially implanted in 3dpf *Tg(fli1a:eGFP)^y1^* larvae. Live confocal imaging of animals at 72 hours post-injection reveals GBM growth and development in the larval zebrafish brain. Scale bar= 50µm.

### Tumor microenvironment affects perivascular glioma invasion in vivo

Cancer arises when normal cells within an organ become misprogrammed. Prior studies have injected non-CNS cancer cells into the periphery of the zebrafish to assess tumor associations with blood vessels ^22, 23^. Therefore, it is logical to inject tumor cells in a tissue in which those cells naturally thrive. To validate the importance of the microenvironment in our perivascular GBM invasion model, we injected D54-MG-tdTomato cells in the trunk near the dorsal fin of 3dpf *Tg(fli1a:eGFP)^y1^;casper* animals. Surprisingly, these glioma cells not only survived, but they radically moved rostrally away from the dorsal fin area and towards the brain by 7dpf (**Figure 4A-C**). These cells extended long projections during invasion (**Supplemental Movie 1),** and unlike when in the brain, these cells did not attach to non-lymphatic, pre-existing blood vessels. This data suggests there are unique trophic factors within the brain environment that attracts glioma cells to the zebrafish vascular network. Interestingly, we observed instances of tumor-induced angiogenesis in 60% of injected animals (n=5 animals), a biology witnessed in other zebrafish cancer models (**Figure 4E,F**) ^22, 24^. These experiments highlight the importance of the microenvironment when modeling and quantifying glioma-vessel interactions.

**Figure 4.**
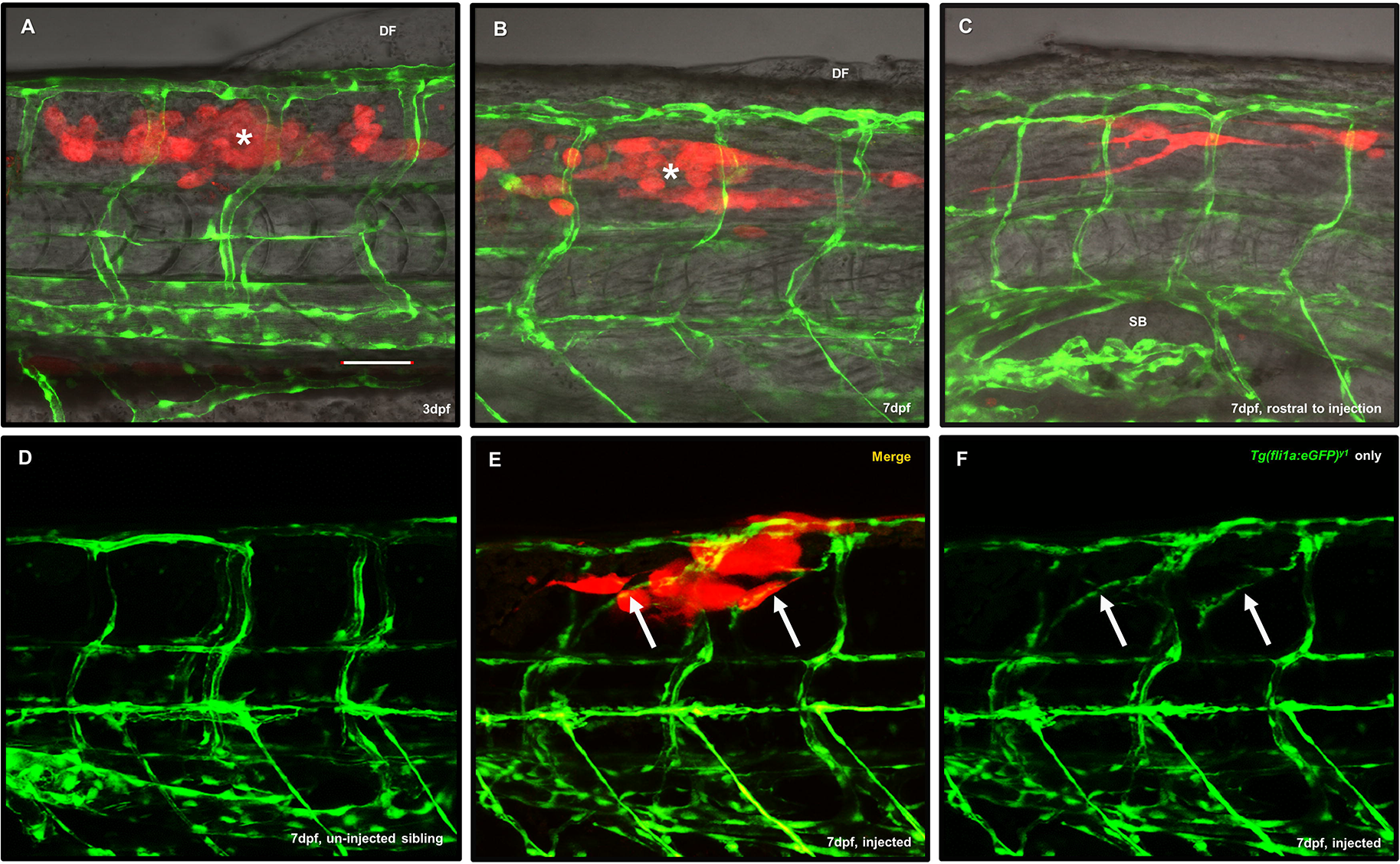
Glioma cells do not migrate along pre-existing peripheral vessels, but do evoke non-CNS angiogenesis. (A) Representative maximum intensity projection image of a 3dpf *Tg(fli1a:eGFP)^y1^;casper* animals (green) injected with 25-50 D54-MGtdTomato cells (red) into the trunk of the muscle (white asterisk) close to the dorsal fin (DF) to survey whether GBM would interact with non-CNS vasculature. (B) The same animal was imaged at 7dpf to survey tumor cell survival and movement from the point of injection (white asterisk). (C) Rostral from the point of injection in the same animal from (B), tumor cells moved down the trunk towards the brain and appear near the swim bladder (SB). *n*=9 animals. (D) A representative image of trunk intersegmental vessel patterning in a 7dpf un-injected *Tg(fli1a:eGFP)^y1^;casper* sibling. (E) We noted instances where tumor cells (red) reaching outward also ensheathed new blood vessels (E,F arrows) that were created after tumor implantation, not seen in un-injected sibling controls. Scale bar=50µm.

### Gross vessel morphology and glut1 expression is not altered after tumor invasion

After determining that tumor cells failed to attach to pre-existing, non-CNS vasculature, we wanted to further investigate other parameters post-invasion. Due to the reports of vascular tone alterations and BBB disruption in both a GBM mouse model and in patient tissue, we assessed the zebrafish brain vascular morphology after 4 days of glioma invasion ^11, 25^. We first asked whether vessel volume was changed in our zebrafish model of perivascular glioma invasion. We utilized volumetric analyses of vessels without tumor burden and vessels on the contralateral brain region contacted by D54-MG-eGFP tumor cells(**Figure 5A,B**). While tumor cells readily ensheathed vessels in the zebrafish brain, we did not find any difference in vessel volume 4 days post-invasion (**Figure 5C**). As the *Tg(glut1b:mCherry)* line expresses mCherry under the control of a reliable BBB marker, *glut1*, we next analyzed the mCherry fluorescence signal of tumor and non-tumor associated vessels in 7dpf animals ^21^. Data from fluorescence intensity analysis revealed that *glut1* expression did not differ between vessels co-opted by a tumor cell soma and vessels in a similar location on the contralateral side of the brain (**Figure 5D**). These live imaging results suggest that co-opted vessels are not visibly compromised, and the BBB specific marker *glut1* was not altered after this 4 day time frame of tumor cell invasion.

**Figure 5.**
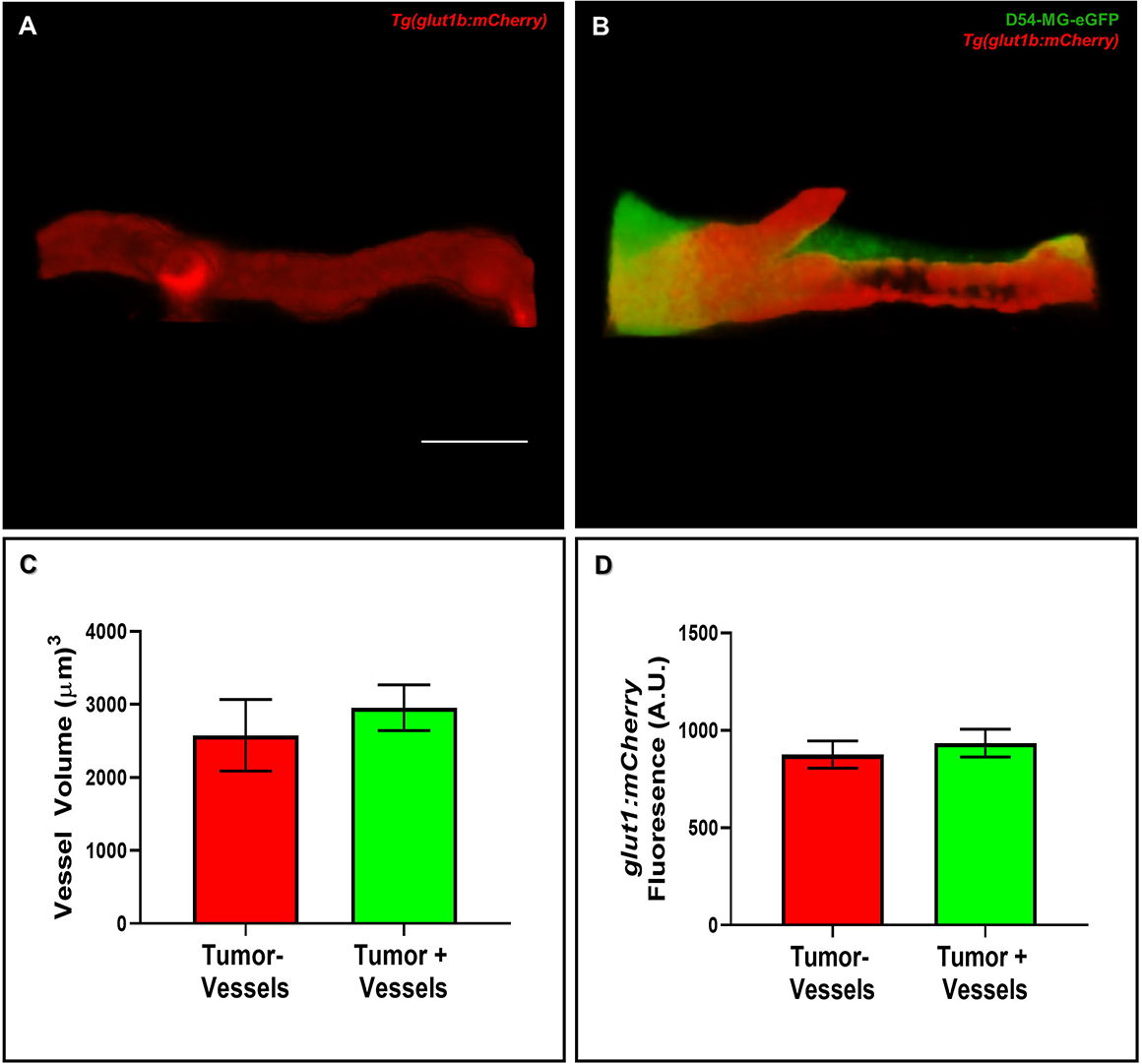
Vessel volume and *glut1* expression is not altered after acute GBM invasion. (A,B) Representative 3D rendered volume views of a blood vessel from the contralateral brain (A) in a region similar to a vessel co-opted by a D54-MG-eGFP tumor cell in a 7dpf zebrafish brain (B). (C,D) The volume and signal intensity of *glut1* in tumor-associated and non-tumor associated vessels were measured in 3D rendered volume views. (C) Quantification of vessel volume. A two-tailed, paired student’s t-test was performed. (D) Quantification of *glut1b:mCherry* fluorescence. A two-tailed, unpaired student’s t-test was performed. Error bars represent mean with standard error of the mean. *n*=8 animals total, 1-3 vessels per animal per group. Scale bar= 10µm.

### Glioma-vessel interactions can be pharmacologically inhibited in vivo

In order to demonstrate whether glioma-vessel interactions could be pharmacologically disrupted *in vivo*, we sought to perform proof-of-principle experiments targeting pathways important for glioma-vessel interactions in GBM mouse models, such as Wnt signaling ^12^. Therefore, we hypothesized that pathways important for mitigating tumor-vessel biology in rodent models would be conserved during perivascular glioma invasion in zebrafish brain ^12^. We hypothesized that adding a small molecule inhibitor of Wnt signaling (XAV939) to the water during tumor invasion, would result in fewer cellular interactions in the zebrafish brain. To test this hypothesis, 4dpf animals were blindly sorted into DMSO vehicle control or XAV939 treatment groups 24h after tumor cell implantation. After 48 hours of bath application, confocal images revealed significantly fewer glioma-vessel interactions (21.57% ± 4.908) in XAV939 treated animals compared to control treated animals (41.5% ± 7.106) (**Figure 6E**). Important to note, the total number of cells was not significantly different after 48 hours of either treatment (**Figure 6F**). While there was a change in cell number after 48 hours of XAV939 treatment (−2.5 ± 6.786, n=6) compared to DMSO vehicle control (11.25 ± 9.013, n=4), this was not statistically significant (**Figure 6G**). These studies suggest that conserved signaling pathways in mammals attract glioma cells to the zebrafish brain vasculature and this *in vivo* system is amenable to chemical perturbation.

**Figure 6.**
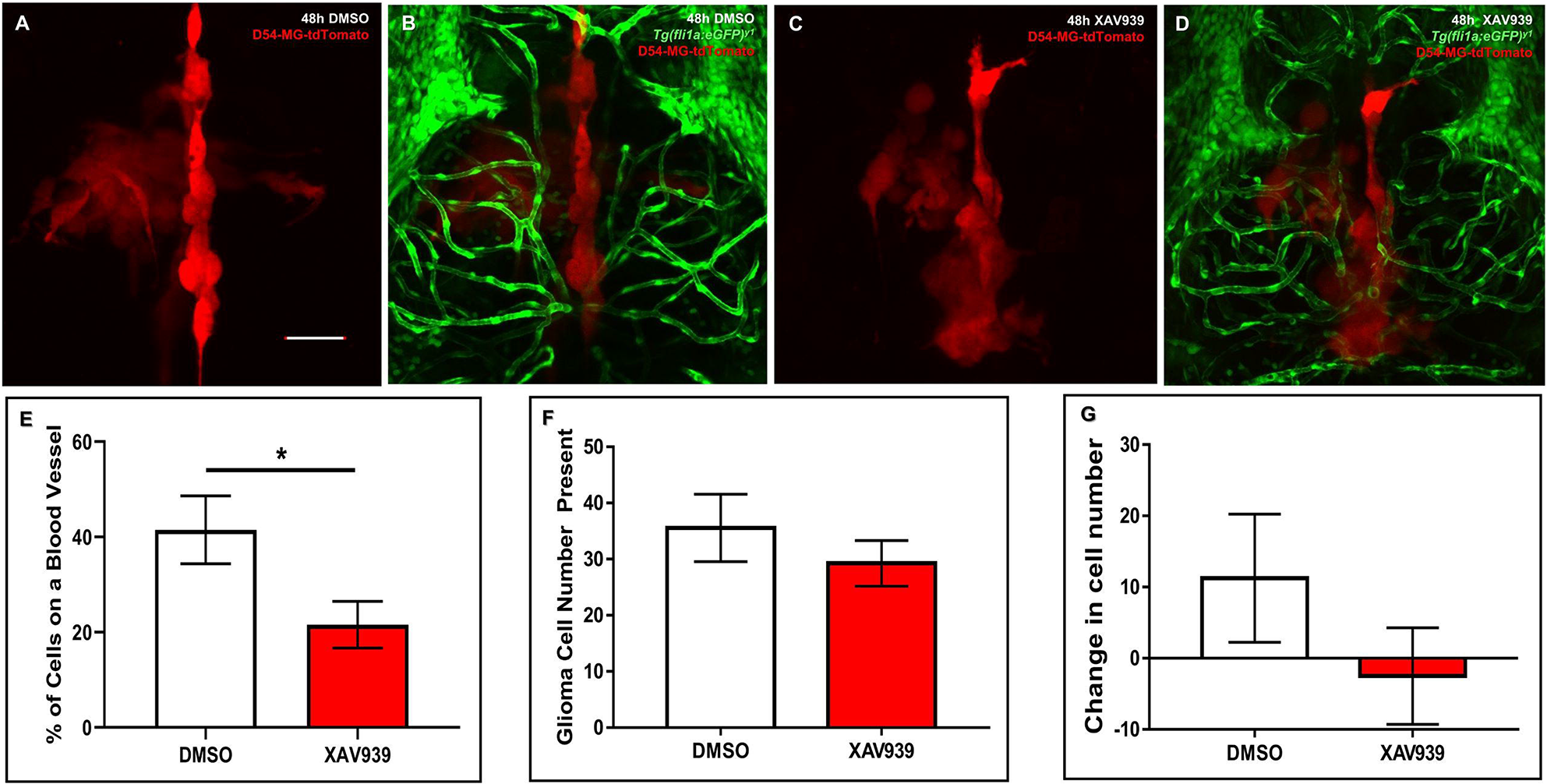
Pharmacological inhibition of Wnt signaling diminishes tumor-vessel interactions. (A-D) Representative maximum intensity projections of tumor burden (red) in vehicle control (A,B) or 30µM XAV939 treated (C,D) animals. Animals were injected with D54-MG-tdTomato cells at 3dpf, unbiasedly sorted, and treated between 4-6dpf. (E) Quantification of tumor-vessel interactions after 48h of vehicle control (DMSO) or 30µM XAV939 treatment. A two-tailed, unpaired t-test was performed (p=<0.05). *n*=6 animals per group. Error bars represent mean with standard error of the mean. (F) Quantification of total glioma cells resulting after 48h of vehicle control (DMSO) or 30µM XAV939 treatment. A two-tailed, unpaired t-test was performed. *n*= 7-8 animals per group. Error bars represent mean with standard error of the mean. (G) Quantification of the change in tumor cell number after 48h of vehicle control (DMSO) or 30µM XAV939 treatment. A two-tailed, unpaired t-test was performed. *n*=4-6 animals per group. Error bars represent mean with standard error of the mean. Scale bar= 50µm.

## Discussion

GBM is a fatal primary brain tumor studied in an extensive array of models. More recently, research groups have implemented xenograft studies to assess glioma invasion in the developing zebrafish. Zebrafish are genetically tractable, share a similar body plan and genetic conservation to mammals, and are optically transparent. The zebrafish does not develop a functional adaptive immune system until 4 weeks of age, and therefore circumvents the need for requiring immunosuppressive chemicals or special animals for injection at early time points ^26^. The adult mouse brain contains roughly 700 times more neuronal cells than the larval zebrafish brain,thus fewer tumor cells need to be injected to visualize similar phenotypes ^27, 28^. These features have presented zebrafish to cancer scientists as an attractive model for dissecting tumor biology and finding new therapeutics ^16, 29^. As the zebrafish model has emerged as a suitable system for dissecting cancer biology, suchstudies have dissected the processes of tumor invasion and angiogenesis ^15, 23^.

However, tumor cells are commonly loaded with lipophilic dyes, which, compared to genetic expression of fluorescent proteins, are not as effective for thoroughly labelling cells and their finer processes for imaging studies ^16, 24^. As it is now widely recognized in the field of GBM biology, gliomas are highly motile tumors known for rapid proliferation and migration. Establishing the biology of perivascular glioma invasion in the zebrafish model would open therapeutic screening possibilities towards a pathological tumor behavior commonly seen in a majority of GBM patients.

Our injection studies reveal the active process of perivascular glioma invasion along the zebrafish CNS vasculature *in vivo*. These observations are exciting for two reasons. First, we demonstrated that important glioma biology could be visualized in zebrafish with only 25-50 cells in 4 days. These experiments utilized significantly fewer cells in a fraction of the time compared to traditional mouse models ^11, 12^. Secondly, a conserved biology must support the attraction of human cells to the developing zebrafish brain vascular network. One of the most critical elements in establishing a model of perivascular glioma invasion is cell number. Other research groups inject widely varied numbers of tumor cells, ranging from fifty to hundreds of tumor cells. We found that fewer cells injected allowed the tumors to migrate. While an individual glioma cell can shrink up to 30% of its volume to invade the mammalian brain, injecting fewer cells likely provides the glioma cells with more room to not only grow but also invade in a smaller space ^18^. By injecting only 25-50 cells, we observed glioma cells associating with the zebrafish brain vasculature within 24 hours post-implantation. This denotes a salient feature of GBM biology; tumor cell attachment to secondary structures during invasion. We also witnessed brain necrosis in animals injected with >50 cells, as well as animals succumbing to their tumor burden (data not shown). Most interestingly, perivascular glioma invasion increased over time, from 3-7dpf. After only 4 days of growth, we witnessed a similar percentage of tumor-vessel interactions as previously demonstrated in a GBM mouse model that also utilized implantation of D54-MG cells ^11^. Furthermore, we observed tumor-vessel interactions with adherent cell lines and patient-derived xenolines. These experiments highlight the model’s accessibility due to what resources may be available to a research group, as PDX lines are time and resource intensive to maintain. Of note, our study is one of the few to utilize pediatric xenoline cells and demonstrate that tumor cell-vessel interactions arise in the context of pediatric CNS cancer *in vivo*.

The extracellular milieu plays a major role during tissue development and for cellular interactions. The unique composition of the brain space is also remodeled in the context of brain tumor progression^30^. When assessing the possibility of perivascular glioma invasion in the periphery of the zebrafish, we did not observe glioma-vessel interactions with the pre-existing vasculature (**Figure 4**). While others have reported glioma induced-angiogenesis after tumor implantation in the zebrafish brain, little tumor-induced angiogenesis occurred after intracranial injections, possibly due to the lack of large tumors that require more nutrients or the presence of vascular endothelial growth factor (VEGF) already in the developing zebrafish brain during active CNS angiogenesis ^16, 31^. Moreover, with distinct extracellular matrix features, such as the selective BBB and a functional lymphatic system, the brain creates an exclusive environment for cancer growth^32^.

While it is common for other cancers to metastasize into the brain, it is possible that specific tropism to the BBB and the surrounding microenvironment prevents CNS tumors from hematagenously spreading into the periphery. For example, studies have demonstrated how astrocytes produce connexin 43 mediated tumor-stromal interactions and how neuronal activity promotes cancer cell proliferation through neuroligin-3 upregulation ^33, 34^. Glioma cells are known to release VEGF during tumor growth ^35^. Because the intersegmental vessels (ISVs) are already patterned by the time we injected cells, it is possible new vessels formed due to tumor cell secreted VEGF. Researchers have witnessed angiogenesis towards non-CNS tumor cells implanted in the zebrafish, a common process that provides a tumor mass with nutrients necessary to maintain uncontrolled cell growth ^23^. Furthermore, in the context of CNS tumors, the BBB vasculature is specialized under a different set of signals than the vasculature within the periphery in both rodents and zebrafish ^36–39^. Interestingly, recent studies injecting non-CNS tumor cells into circulation of the developing zebrafish demonstrated these cancers attached to ISVs unlike our brain tumor cells ^40^. Therefore, the process of angiogenesis (the formation of new blood vessels) and perivascular invasion (attraction of cells to pre-existing vessels for cell migration) underscores different biology in GBM. While these studies support the examination of brain tumors within their tissue of origin, the zebrafish could provide an interesting model to dissect brain tumor preferences for the BBB versus peripheral vasculature attachment. Additionally, such experiments would be easily achievable with the zebrafish model system.

As tumor cells secrete a myriad of destructive molecules during invasion, a theoretical side effect of perivascular glioma invasion is disruption of the vasculature’s integrity ^41^. GBM models and morphological analysis of patient tissue have demonstrated a change in vessel response and BBB breach via the loss of tight junction proteins ^11, 25^. While the BBB was not a direct focus of this study, we looked at a few potential effects of perivascular glioma invasion on the zebrafish CNS vasculature. Zebrafish possess the chemical and physical barriers necessary to make a functional BBB early in development, also making it easy to survey in our model of perivascular glioma invasion ^21, 42^. We found that vessel volume and BBB integrity via expression of *glut1* was not altered in our zebrafish model of perivascular glioma invasion. Our *glut1* data aligns with a study of xenograft models and patient tissue demonstrating normal expression of GLUT1 at vessels co-opted by tumor cells ^43^. While previous studies showed a decrease in tight junction protein expression, it would be an interesting focus to use this zebrafish model to investigate if changes in BBB proteins occur before or after BBB dysfunction during perivascular tumor invasion.

One of the ultimate goals of establishing a zebrafish model of perivascular glioma invasion was to demonstrate the feasibility of using small molecules that could target mechanisms necessary for glioma-vessel interactions. The zebrafish is an ideal organism for *in vivo* cancer drug discovery, as animals can be successfully xenografted with tumors in a short time span, and are amenable to high-throughput chemical assays ^44^. In fact, zebrafish “avatars” are becoming successful platforms for drugs to make it into clinical trials ^45^. In the field of GBM biology, most studies assess overall tumor cell growth, but few have investigated means for the chemical disruption of glioma-vessel communication. Previous research has begun to dissect pathways involved in perivascular glioma invasion mouse models, specifically through Wnt and Bradykinin signaling ^12, 46^. Zebrafish express homologous proteins for both the Wnt and Bradykinin pathways, but the function of these homologs in zebrafish cancer models remain unexplored ^47, 48^. We demonstrate that tumor-vessel interactions in the zebrafish brain can be disrupted by the Wnt signaling antagonist XAV939, supporting the potential of BBB-specific cues that mediate perivascular glioma invasion ^37^. Wnt signaling inhibition was not novel in the context of glioma biology but was in regards to our zebrafish model of tumor invasion. As the Wnt/beta-catenin signaling cascade is targeted in clinical trials, the use of XAV939 in our zebrafish tumor xenograft model highlights the suitability of the zebrafish in the preclinical pipeline ^49^. These proof-of-principle small molecule experiments demonstrate the practicality of using this zebrafish model to identify compounds of interest that can target processes important to glioma invasion. Determining the underlying behavior that supports tumor growth would cultivate novel therapies, as migratory cell populations evade the current GBM standard of care. Therefore, this established zebrafish model of perivascular glioma invasion generates an additional platform for dissecting tumor cell biology and progressing cancer drug discovery.

## Supporting Information

Glioma invasion in a non-specific pathogen free transgenic line, *Tg(glut1b:mCherry)*; Lipophilic dye labeled glioma xenoline implanted into the *Tg(fli1a:eGFP)^y1^* zebrafish brain; 3D-rendered movie of glioma cells in the periphery of a 10 days post-fertilization *Tg(fli1a:eGFP)^y1^* zebrafish larvae.

## Acknowledgements

The authors would like to acknowledge and thank Miss Joelle Martin and AnnaLin Woo for constructive criticism of the manuscript, Mr. Tré Mills for assistance with the volumetric analysis protocol, Dr. Emily Thompson for transducing the D2159MG xenoline for mCherry expression, the Pan laboratory for use of their stereoscope and intellectual discussion, and Dr. Michael Taylor for use of the *Tg(glut1b:mCherry)* line.

---

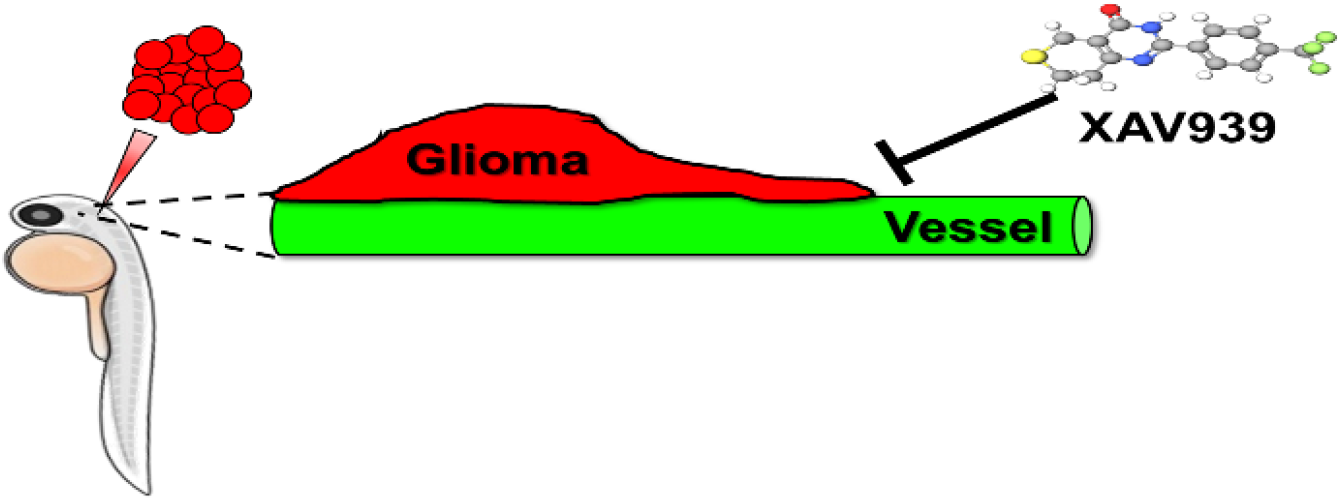
Graphic synopsis: Glioma is a highly invasive brain cancer, as a result of tumor cells migrating through the brain via single cell vessel co-option. While rodents are traditionally used to study tumor biology, zebrafish possess many advantages such as ease of live imaging and chemical screening. We demonstrated and characterized the suitability of utilizing a zebrafish xenograft model to study perivascular glioma invasion. Furthermore, we identified that the Wnt signaling inhibitor, XAV939, inhibits perivascular tumor interactions in zebrafish brain. These studies demonstrate the feasibility of utilizing zebrafish as a platform for future targets in tumor drug discovery.

## Notes

### Competing Interest Statement

The authors have declared no competing interest.

### Summary of Updates

Figures 1 and 6 revised. Results updated to clarify the importance of cell number injected. Discussion updated to clarify importance of extracellular matrix environment and use of XAV939.

