## Supplementary figures and images for "Fishing for contact: Modeling perivascular glioma invasion in the zebrafish brain"

### Supplemental Figure 1

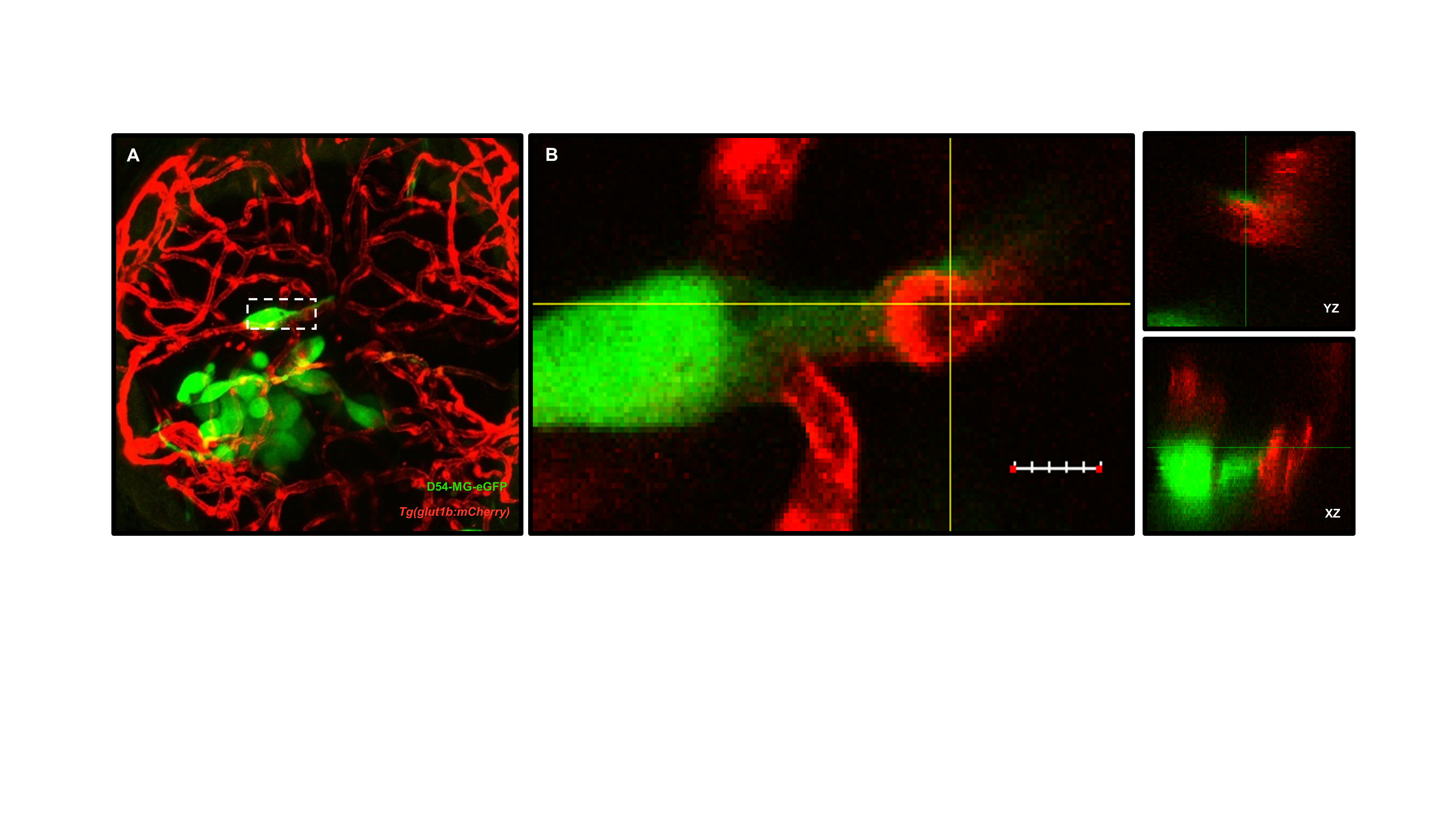

### Supplemental Figure 2

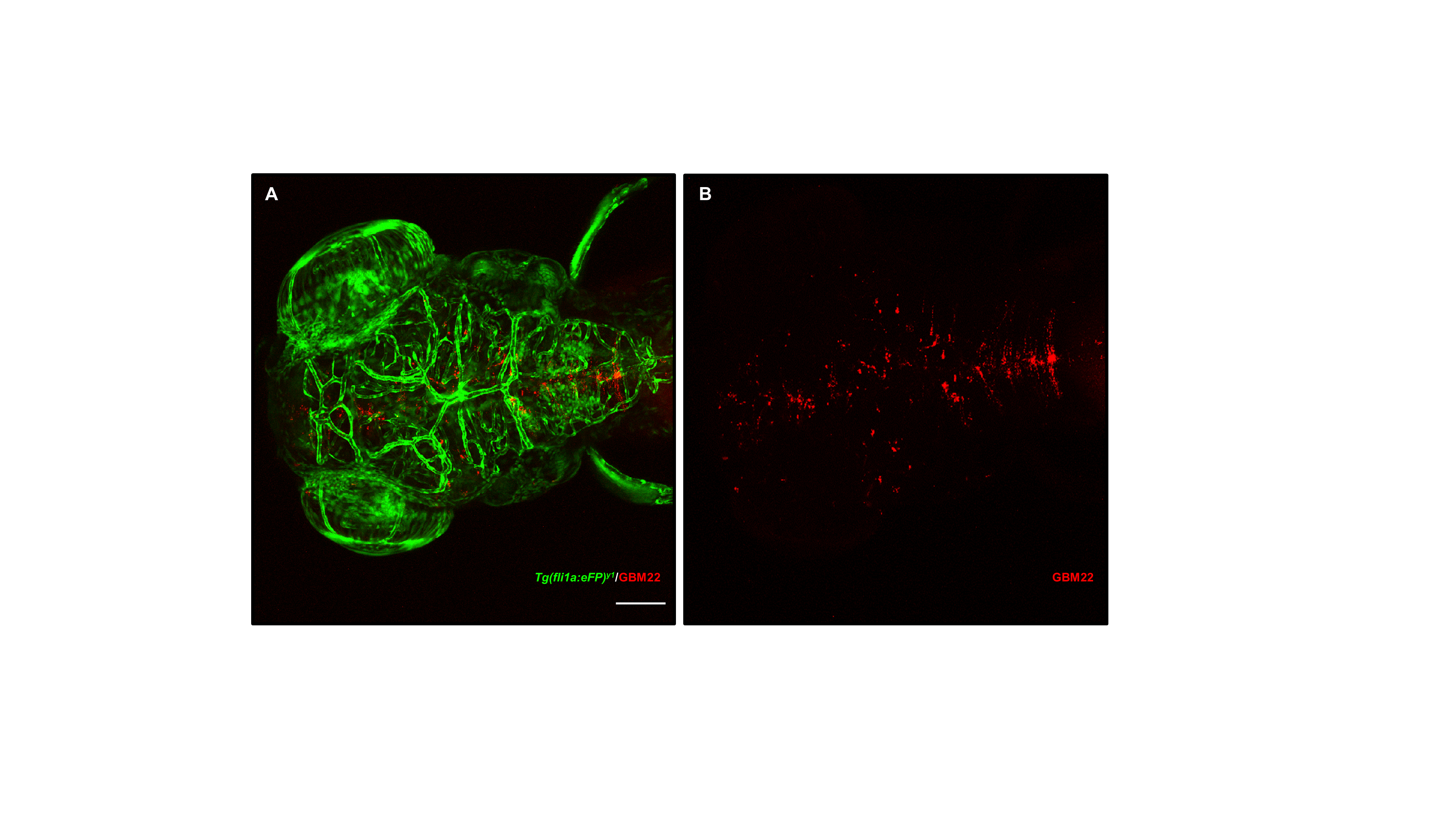
