## Supplemental Figure legends for "Fishing for contact: Modeling perivascular glioma invasion in the zebrafish brain"

**Table of Contents:**

| Data | Page Number |
| --- | --- |
| Figure 1 | 6 |
| Figure 2 | 7 |
| Figure 3 | 7 |
| Figure 4 | 7 |
| Figure 5 | 8 |
| Figure 6 | 8 |
| Figure S-1 | 6 |
| Figure S-2 | 7 |
| Movie S-1 | 7 |

**Figure S-1. Tumor-vessel interactions are maintained in non-specific pathogen free zebrafish lines.** (A) A maximum intensity projection of a 7dpf *Tg(glut1:mCherry)* larvae (red) implanted with D54-MG-eGFP tumor cells (green). (B) An area (dotted white box in A) of a representative Z-plane from the confocal stack showing the corresponding orthogonal planes, between the vessel (red) and glioma cell (green) signals. *n*=18 animals. Scale bar= 10µm.

**Figure S-2.** **Labeling glioma cells with lipophilic dyes is not ideal for visualizing perivascular glioma invasion.** (A) A maximum intensity projection of the whole brain of a 4dpf *Tg(fli1a:eGFP)y1;casper* (green) larvae with GBM22 cells (red) loaded with a lipophilic dye 24 hours post-injection. (B) A maximum intensity projection image of the red channel displaying the punctate staining of the human glioma cells. Scale bar= 100µm.

**Movie S-1. Human glioma cells expand within in the periphery of the developing zebrafish larvae.** A 3D rendered volume view movie of a 10dpf *Tg(fli1a:eGFP)^y1^;casper* zebrafish, 1 week post tumor implantation. Note that the tumor cells (red) do not interact with any pre-existing vasculature (green) and have migrated towards the rostral end of the animal, from the dorsal fin area past the swim bladder.
